# Ultrasound-Activated Nanobubbles Induce Durable Systemic Antitumor Immunity

**DOI:** 10.64898/2026.03.25.714247

**Authors:** Anubhuti Bhalotia, Pinunta Nittayacharn, Diarmuid W. Hutchinson, Andrew Cheplyansky, Koki H. Takizawa, Abraham Nidhiry, Shreya Hariharan, Ashley Novak, Arya Iyer, Meghna Mehta, Theresa Kosmides, Reshani Perera, Inga M. Hwang, Agata A. Exner, Efstathios Karathanasis

**Affiliations:** Department of Biomedical Engineering, School of Medicine, Case Western Reserve University, Cleveland, Ohio 44106, USA; Department of Biomedical Engineering, Faculty of Engineering, Mahidol University, Phuttamonton, Nakorn Pathom, 73170, Thailand; Department of Radiology, School of Medicine, Case Western Reserve University, Cleveland, Ohio 44106, USA; Case Comprehensive Cancer Center, School of Medicine, Case Western Reserve University, Cleveland, Ohio 44106, USA

**Author notes:** Co-corresponding authors: Efstathios Karathanasis and Agata Exner. **Data availability statement:** The data supporting the findings of this study are available within the article and its supplementary material. **Approval statement:** All animal procedures were conducted under a protocol (protocol number: 2016-0115) approved by the Institutional Animal Care and Use Committee (IACUC) of Case Western Reserve University (CWRU).

**Keywords:** ultrasound, nanobubbles, immunotherapy, mechanical immunomodulation, breast cancer

## Abstract

Clinical outcomes in aggressive breast cancer vary widely, in part because the tumor microenvironment is structured to exclude immune infiltration. Low antigen load, dysfunctional antigen-presenting cells, T cell exclusion and exhaustion, and a stiff extracellular matrix that physically restricts immune cell trafficking work together to form a suppressive barrier that current immunotherapies struggle to overcome. We addressed this barrier using ultrasound (US)-activated nanobubbles (NBs), a drug-free intervention based on perfluoropropane-filled nanoparticles. The size and deformable phospholipid shell enable NBs to achieve deep tumor penetration and a uniform distribution throughout the entire tumor. Upon ultrasound activation, NBs generate localized mechanical forces that restore extracellular matrix elasticity, disrupt tumor transport barriers, and drive HMGB1 release, re-engaging endogenous antitumor immunity without pharmacological agents. In a syngeneic triple-negative breast cancer model, US-NB treatment depleted immunosuppressive myeloid cells 3-fold within 3 hours, followed by a greater than 5-fold increase in the ratio of antigen-experienced to suppressive T cells at 48 hours. US-NB drives rapid infiltration of CD4^+^ and CD8^+^ T cells within 48 hours. US-NB treatment achieved an 85% cure rate in the D2A1 model; cured animals maintained durable systemic immune memory, rejecting both local and systemic tumor rechallenge. Consistent therapeutic benefit was observed in a luminal B-like mammary tumor model (E0771), supporting activity across breast cancer subtypes. These results establish US-NB mechanical immunomodulation as a drug-free therapeutic strategy capable of generating robust and durable antitumor immunity, acting through biophysical tissue properties rather than tumor-specific molecular targets.

**GRAPHICAL ABSTRACT:** 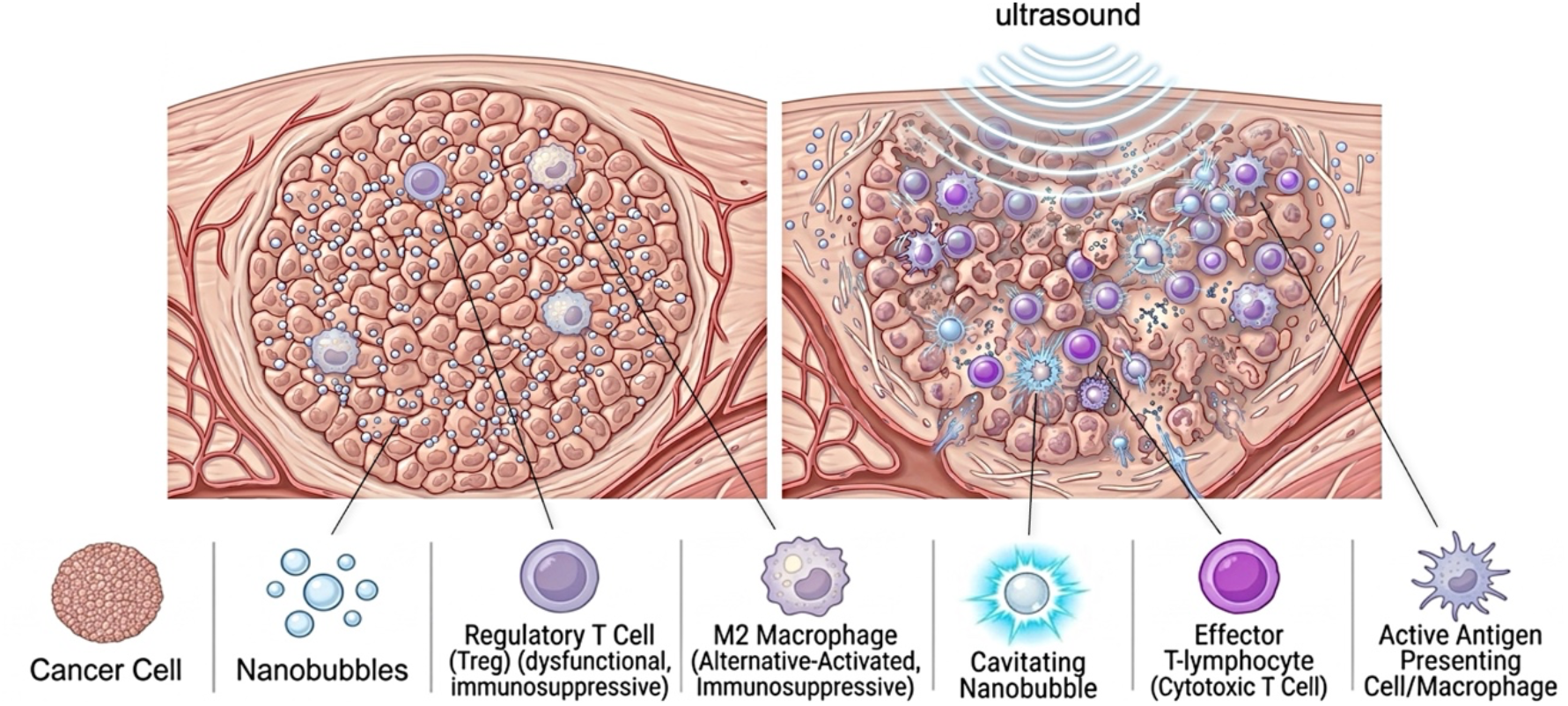

## INTRODUCTION

Breast cancer is among the most biologically heterogeneous solid malignancies, with clonal diversity generating resistant subpopulations that evade targeted and cytotoxic therapies across subtypes.^1^ The tumor immune microenvironment drives this therapeutic resistance through limited antigen availability, dysfunctional antigen-presenting cells, T-cell exclusion, and a dense, stiff extracellular matrix (ECM) that restricts immune cell trafficking.^2-5^ As a result, current immunotherapies (*e*.*g*., immune checkpoint inhibitors) are largely ineffective.^5,6^

Effective treatment therefore requires a strategy that does not depend on specific tumor molecular features while simultaneously restoring immune access to the tumor and priming durable, systemic antitumor immunity.^7^ In situ vaccination strategies address this need by converting the tumor into a source of endogenous antigens that educate the immune system without prior molecular characterization.^8,9^ Physical ablation modalities, notably high-intensity focused ultrasound, have been explored in this context, releasing damage-associated molecular patterns (DAMPs) into the TME.^10-12^ However, thermal approaches (e.g., HIFU) denature proteins through coagulative necrosis, compromising antigen integrity,^13,14^ while non-thermal approaches such as histotripsy use extreme peak negative pressures (∼25 MPa) to fractionate the entire tumor into acellular debris, eliminating the in situ tissue architecture that supports antigen presentation and immune cell trafficking.^15,16^ Although tumor destruction is achieved in both cases, the resulting loss of this architecture impairs productive immune engagement, yielding responses that are typically transient and insufficient to generate durable protective memory without co-administration of immunotherapeutic agents. These issues highlight mechanical, rather than disruption as a critical and underexplored mechanism for effective in situ immunomodulation.

Ultrasound-activated (US) nanobubbles (NBs) offer a mechanical, non-pharmacological strategy for in situ immunomodulation.^17,18^ NBs are echogenic, perfluoropropane gas-filled particles with compressible gas cores and robust yet deformable phospholipid shells.^19-21^ Contrary to larger bubbles (e.g., microbubbles), NBs are able to penetrate deep into tissue and efficiently distribute throughout the entire tumor mass, creating widespread reservoirs of nanoscale gas bubbles.^17^ This is a unique feature of NBs as a result of their deformable shell, nanoscale size and compressible gas core. Mild US causes the NBs to oscillate and cavitate, which generates localized, gentle mechanical forces in the surrounding tissues capable of disrupting the tumor transport barriers. The generated microstreaming effectively breaks down the dense tumor tissue ECM (e.g., collagen) and restores the tumor tissue elasticity without eliminating key immune cell populations.^17^

We hypothesize that US-NB-mediated mechanical disruption of the tumor ECM is sufficient to reverse local immunosuppression and generate durable systemic antitumor immunity without pharmacological intervention. To test this, we evaluated US-NB as a standalone intervention in aggressive syngeneic breast cancer models. Through time-resolved immunophenotyping, we observed that a single US-NB treatment depleted monocytic myeloid-derived suppressor cells (mMDSCs) 3-fold within 3 hours, followed by macrophage polarization toward an antigen-presenting phenotype and a greater than 5-fold shift in the ratio of antigen-experienced to suppressive T cells by 48 hours. This coordinated immune reprogramming extended to the tumor-draining lymph node and spleen, where antigen-experienced CD44^+^ T cell clones expanded across both compartments. Therapeutically, US-NB achieved an 85% cure rate in established D2A1 tumors, and cured animals rejected both local and systemic tumor rechallenge months later, confirming durable immune memory. Consistent therapeutic benefit was observed in the E0771 luminal B-like mammary model, supporting activity across breast cancer subtypes. These data demonstrate that US-NB mechanical immunomodulation is sufficient to drive a curative antitumor immune response without pharmacological intervention.

## RESULTS

### Nanobubbles are sequestered within tumors with minimal systemic leakage

Effective mechanical disruption of the tumor mass requires that NBs distribute throughout the entire tumor volume, maximizing cavitation coverage while minimizing systemic exposure and off-target tissue damage. We first assessed the spatiotemporal distribution of intratumorally injected nanobubbles using a dual-modality imaging approach. Real-time contrast-enhanced ultrasound (CEUS) imaging provided dynamic kinetic information at multiple timepoints, while ex vivo fluorescence imaging enabled quantitative organ-level biodistribution analysis. We performed CEUS imaging at 18 MHz across the tumor, kidney, and liver at 0-, 90-, and 180-minutes post-injection, sampling three imaging planes per organ in three mice (Fig. 1A). At the 3-hour endpoint, we harvested the tumor, liver, lungs, kidney, spleen, heart, tumor-draining lymph node, and plasma, then quantified fluorescence of Cy5-labeled nanobubbles using IVIS imaging with analysis of average radiant efficiency for each region of interest. This enabled direct kinetic correlation between real-time contrast dynamics (CEUS) and terminal organ-level nanobubble distribution (IVIS imaging).

**Figure 1.**
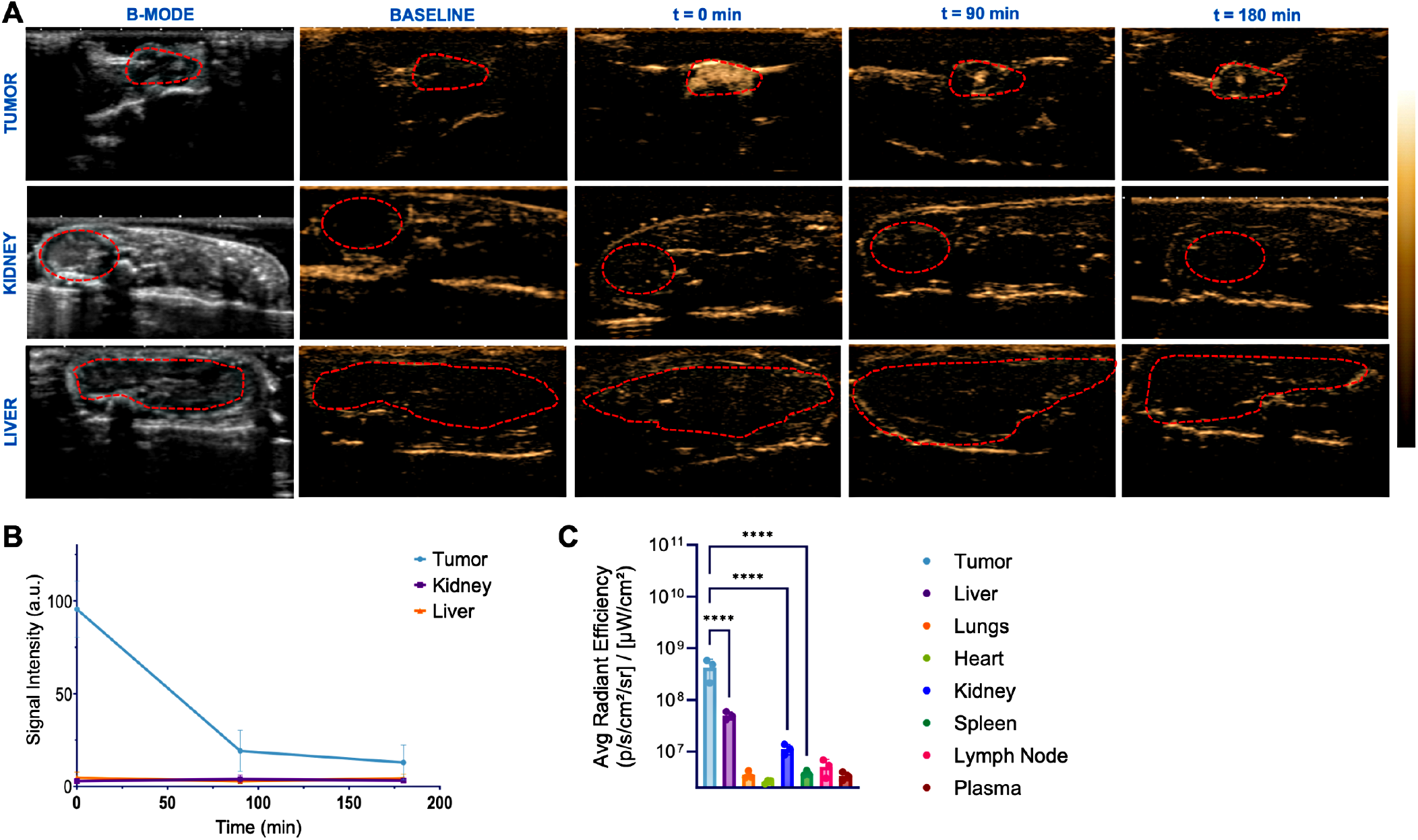
Nanobubble retention and biodistribution characterization. D2A1 tumor-bearing mice received intratumoral injection of Cy5-labeled NBs and were assessed over 3 hours. (A) Representative CEUS imaging frames of tumor, kidney, and liver at 0-, 90-, and 180-minutes post-injection. (B) Nonlinear contrast signal quantified from CEUS imaging averaged across 3 imaging planes per organ (n=3 mice), showing progressive tumor signal decay over 3 hours. (C) Ex vivo biodistribution of Cy5-labeled NBs in major organs at 3 hours post-injection, reported as average radiant efficiency [p/s/cm^2^/sr]/[µW/cm^2^] per IVIS region of interest. Statistics: one-way ANOVA with Tukey’s post-hoc test (n=3 mice; mean ± SD).

The characterization of Cy5-conjugated NBs is shown in Supplementary Fig. S1. The labeled NBs had a hydrodynamic diameter ranging from 200-400 nm and a buoyant (gas-filled) NB concentration of 4.7 x 10^11^ particles/mL, consistent with the unlabeled standard NB formulation previously reported.^17^ Consistent with prior work,^17^ CEUS imaging confirmed rapid and complete tumor filling immediately post-injection (Fig. 1A). NB shell deformability allows passive compression and penetration through the dense tumor ECM, distributing NBs throughout the tumor volume rather than pooling near the injection tract. More than 85% of the initial contrast signal cleared within 3 hours (Fig. 1B), consistent with gas diffusion-driven loss of echogenicity as perfluoropropane dissipates from the NB cores. No signal above background was detected in the kidney or liver at any timepoint, confirming that intratumorally injected NBs do not enter systemic circulation.

Ex vivo quantification of Cy5-labeled NBs at the 3-hour endpoint confirmed the minimal systemic leakage. Tumor tissue retained the highest radiant efficiency (Fig. 1C), establishing the baseline for comparison. The liver exhibited a modest accumulation of NBs with an average radiant efficiency of approximately 5x10^7^ p/s/cm^2^/sr]/[µW/cm^2^] per ROI, representing roughly 10-fold lower accumulation than tumor tissue. The kidney exhibited substantially lower fluorescence, with radiant efficiency approximately 40-fold lower than the tumor. In contrast, all other organs (lungs, heart, spleen, tumor-draining lymph node, and plasma) demonstrated radiant efficiency values below 10^7^, effectively background-level fluorescence relative to tumor signal.

This biodistribution data demonstrates that intratumorally injected NBs efficiently infiltrate and remain confined within tumor tissue with minimal systemic dissemination. The combination of immediate tumor filling, enabled by NB deformability, and intratumoral gas diffusion-driven signal loss establishes a favorable pharmacokinetic profile for localized intratumoral therapy. This selective tumor sequestration substantially mitigates the risk of off-target cavitation-induced tissue damage upon ultrasound activation.

### US-NB reprograms the innate immune landscape toward a pro-inflammatory, antigen-presenting phenotype

Having confirmed that NBs distribute uniformly throughout the tumor and remain confined without systemic leakage, we next asked whether ultrasound-activated cavitation was sufficient to reprogram the immunosuppressive tumor microenvironment. Using flow cytometry, we assessed innate immune changes at 3 and 48 hours following a single US-NB treatment These two timepoints were selected to distinguish the acute mechanical response from sustained immune reprogramming. The flow gating strategy is shown in Supplementary Fig. S2.

US-NB treatment was initiated when tumors reached approximately 50 mm^3^. NBs were diluted 1:5 in PBS and approximately one billion NBs were administered intratumorally. Therapeutic ultrasound was applied immediately using an unfocused transducer with a 1 cm^2^ effective radiating area at 3.3 MHz, 2.2 W/cm^2^, and 50% duty cycle for 10 minutes with acoustic coupling gel, building on our previously developed methods.^17^ The 10-minute treatment duration was selected to deliver sufficient cumulative cavitation energy for standalone immunomodulation without a co-administered drug. Core acoustic parameters (frequency, intensity, duty cycle, transducer geometry, and PNP) matched those validated in our prior work, where US-only controls confirmed that these settings do not independently produce tissue remodeling or immune changes in the absence of NB-mediated cavitation.^17^ Infrared thermometry confirmed tumor surface temperature of 36.8°C after treatment, and all mice recovered immediately upon removal from isoflurane, with no observable effect on health.

US-NB treatment caused rapid depletion of immunosuppressive myeloid populations. Monocytic MDSCs (mMDSCs; CD11b^+^Ly6C^+^Ly6G^−^) were reduced approximately 3-fold as early as 3 hours post-treatment, and this reduction was sustained at 48 hours (Fig. 2A). Simultaneous with mMDSC depletion, the macrophage compartment polarized toward a pro-inflammatory, antigen-presenting phenotype. Antigen-presenting macrophages (F4/80^+^MHC-II^+^) expanded 1.2-fold at 3 hours and remained elevated at 1.4-fold by 48 hours (Fig. 2B). M1-like macrophages (F4/80^+^MHC-II^+^CD86^+^), indicative of Th1 polarization, doubled in proportion by 48 hours (Fig. 2C). The M1-to-M2 macrophage ratio, calculated as CD86^+^ to CD206^+^ frequency, rose from 1.5-fold at 3 hours to 1.8-fold by 48 hours (Fig. 2D). Elevated M1-to-M2 ratios associate with favorable clinical outcomes in breast cancer, reflecting a microenvironment that supports rather than suppresses immune cell activation.^22,23^ Together, rapid mMDSC depletion and sustained macrophage polarization toward an antigen-presenting M1 phenotype established a pro-inflammatory tumor microenvironment that opposes immunosuppression and promotes immune cell infiltration.

**Figure 2.**
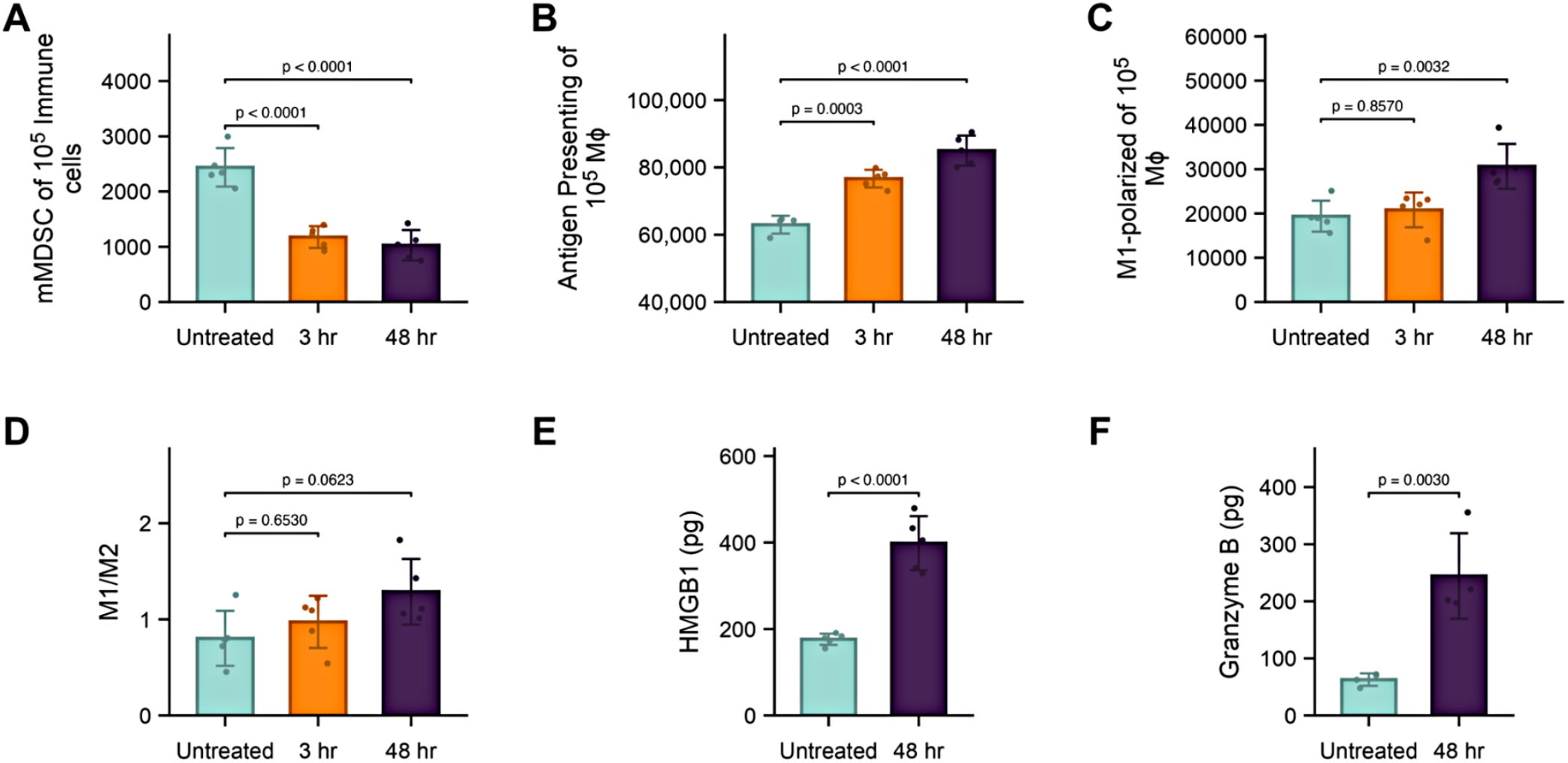
Innate immune reprogramming in the tumor microenvironment following US-NB treatment. D2A1 tumor-bearing mice received a single US-NB treatment and tumor immune infiltrates were assessed by flow cytometry at 3- and 48-hours post-treatment. (A) Monocytic MDSCs (CD11b^+^Ly6C^+^Ly6G^−^). (B) Antigen-presenting macrophages (F4/80^+^MHC-II^+^). (C) M1-like macrophages (F4/80^+^MHC-II^+^CD86^+^). (D) M1-to-M2 macrophage ratio (CD86^+^ to CD206^+^). Tumor homogenates at 48 hours: (E) HMGB1 and (F) Granzyme B quantified by ELISA. Statistics: one-way ANOVA with Tukey’s post-hoc test for flow cytometry data; unpaired t-test for ELISA data (n=5 per group; mean ± SD). Gating hierarchies in Supplementary Fig. S2.

To assess whether US-NB-induced innate reprogramming generated markers consistent with immunogenic cell death, we quantified HMGB1 and Granzyme B in tumor homogenates at 48 hours via ELISA. HMGB1, a DAMP released during cell death that drives dendritic cell maturation and T cell priming,^24,25^ was elevated 3-fold relative to untreated controls (Fig. 2E). Granzyme B, a marker of cytotoxic granule release and active effector function, was elevated 5-fold compared to untreated tumors (Fig. 2F). The combined release of these molecules at 48 hours is consistent with active cytotoxic immune engagement within the treated tumor.

Rapid mMDSC depletion, macrophage polarization toward an antigen-presenting phenotype, and concurrent elevation of HMGB1 and Granzyme B within 48 hours of a single US-NB treatment indicate that mechanical disruption alone is sufficient to dismantle the innate suppressive architecture of the tumor microenvironment and generate conditions permissive for adaptive immune engagement.

### US-NB drives rapid T cell infiltration and activation in the tumor

The innate reprogramming established by US-NB treatment was followed by a rapid and substantial reorganization of the intratumoral T cell compartment. Total T cell abundance (CD3ε^+^) increased 2-fold by 48 h, with the infiltrating population composed of both helper and cytotoxic subsets. CD4^+^ T cells increased 9.3-fold relative to total live cells, while CD8^+^ T cells expanded 12.4-fold (Fig. 3A-C). This expansion of both helper and cytotoxic lineages is mechanistically synergistic for tumor control. The speed and magnitude of this expansion, particularly the 12.4-fold CD8^+^ increase within 48 hours, is consistent with a pro-inflammatory microenvironment that both recruits circulating T cells and relieves suppression of tumor-resident populations.

**Figure 3.**
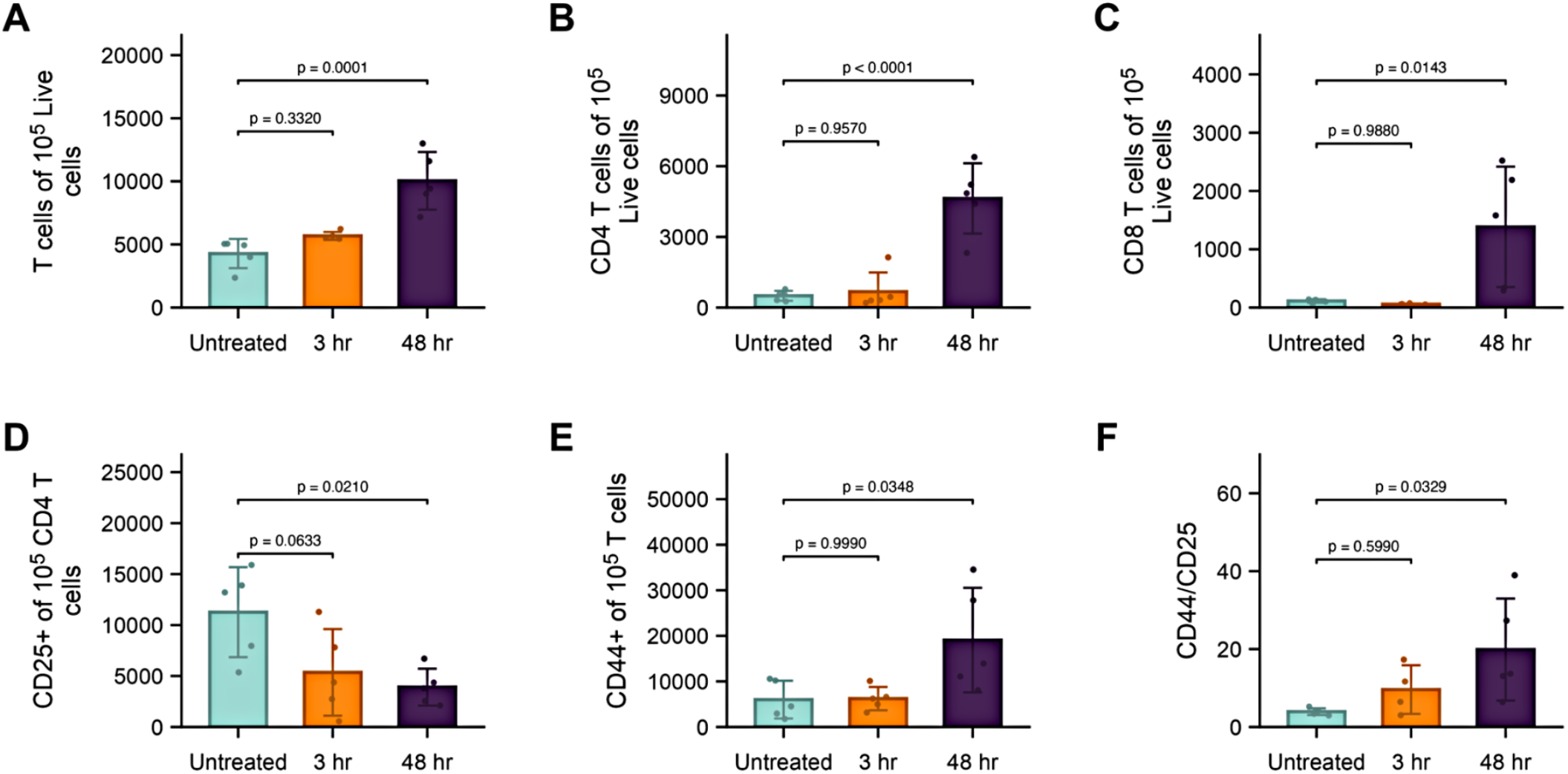
Adaptive T cell infiltration and functional reprogramming in the tumor microenvironment. Flow cytometry of D2A1 tumor single-cell suspensions at 48 hours post-treatment. Frequency of (A) total T cells (CD3ε^+^), (B) helper T cells (CD3ε^+^CD4^+^), and (C) cytotoxic T cells (CD3ε^+^CD8^+^) normalized to live cells. (D) CD4^+^CD25^+^ T cells. (E) CD44^+^ antigen-experienced T cells. (F) CD44^+^ to CD25^+^ T cell ratio. Statistics: unpaired t-test (n=5 per group; mean ± SD). Gating hierarchies in Supplementary Fig. S2.

The infiltrating T cell population demonstrated active antigen-driven engagement. CD4^+^CD25^+^ T cells declined 3-fold between 3 and 48 hours (Fig. 3D). In the tumor microenvironment, CD4^+^CD25^+^ T cells are predominantly regulatory T cells;^26,27^ intratumoral Regulatory T cells accumulation is a well-established mechanism of immune evasion in solid tumors and associates directly with suppression of CD8^+^ effector function and poor clinical outcomes. Their depletion relieved a primary suppressive brake on the effector compartment. Concurrently, CD44^+^ antigen-experienced T cells expanded 3.2-fold (Fig. 3E). CD44 upregulation is a defining marker of T cells that have received activation signals and are committed to effector function and memory differentiation.^28,29^ The CD44^+^ to CD25^+^ ratio shifted from 3.9 in untreated tumors to 19.8 by 48 hours (Fig. 3F), a greater than 5-fold shift from suppressive to effector dominance that reflects a fundamental functional reprogramming of the intratumoral T cell compartment.

### Secondary lymphoid organs undergo robust adaptive immune expansion following US-NB treatment

The intratumoral T cell changes observed at 48 hours raised the question of whether the immune response had propagated systemically. We assessed the spleen and tumor-draining lymph node (TDLN) at 3- and 48-hours post-treatment. As secondary lymphoid organs, TDLNs initiate tumor-specific T cell priming from tumor antigens and the spleen amplifies and shapes systemic antitumor responses.^30,31^ TDLN and spleen can assess whether US-NB links local priming to systemic immunity, including the memory needed to prevent recurrence and metastasis.

NB cavitation initiates the mobilization of innate effector populations to systemic sites beyond the primary tumor. By 48 h post-treatment, antigen-presenting macrophages (F4/80^+^MHC-II^+^) in the spleen expanded 2.0-fold (Fig. 4A), while antigen-presenting dendritic cells (CD11c^+^MHC-II^+^) increased ∼3-fold (Fig. 4B). This innate expansion demonstrates that the immunogenic signals generated by US-NB are not confined to the TME and have spread to distant lymphatic sites such as the spleen. Antigen-presenting macrophages (F4/80^+^MHC-II^+^) in the TDLN expanded 13-fold by 48 h (Fig. 4C). This response was paired with a 14-fold surge in Natural Killer (NK) cells (CD49b^+^; Fig. 4D). This innate expansion in the TDLN created the antigen-presenting and innate effector conditions necessary to support downstream adaptive T cell activation.

**Figure 4.**
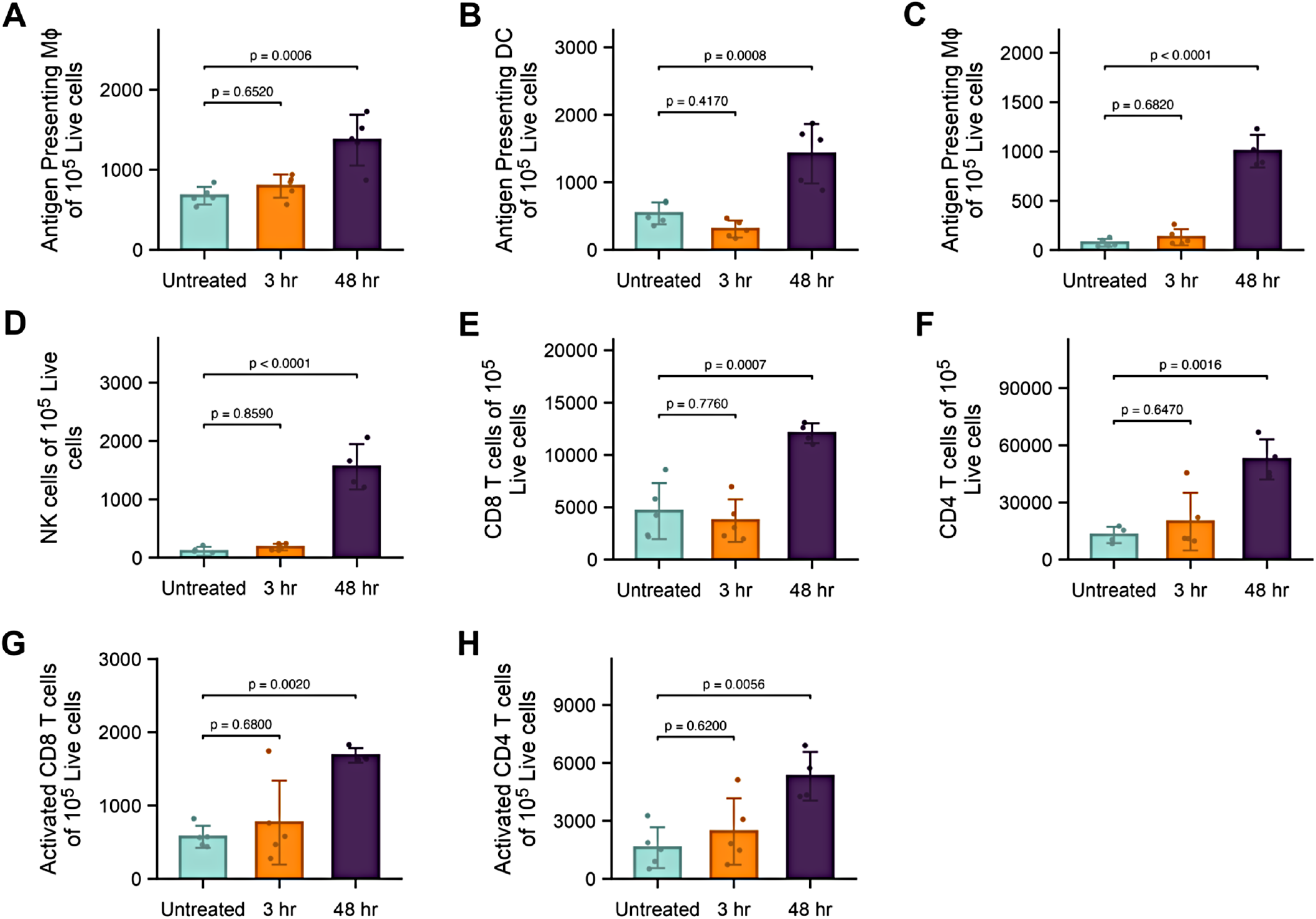
Innate and adaptive immune expansion in secondary lymphoid organs following US-NB treatment. D2A1 tumor-bearing mice received a single US-NB treatment; spleens and TDLNs were harvested at 3- and 48-hours post-treatment. Spleen: (A) antigen-presenting macrophages (F4/80^+^MHC-II^+^); (B) antigen-presenting dendritic cells (CD11c^+^MHC-II^+^). TDLN: (C) antigen-presenting macrophages (F4/80^+^MHC-II^+^); (D) NK cells (CD49b^+^); (E) CD8^+^ T cells; (F) CD4^+^ T cells; (G) CD8^+^CD44^+^ antigen-experienced T cells; (H) CD4^+^CD44^+^ antigen-experienced T cells. Statistics: one-way ANOVA with Tukey’s post-hoc test (n=5 per group; mean ± SD). Gating hierarchies in Supplementary Fig. S2.

Driven by the activation of innate immune cells, the TDLN supported a robust expansion of antigen-specific T cell subsets. We observed a synergistic rise in both CD4^+^ helper (4.1-fold) and CD8^+^ cytotoxic (2.6-fold) lineages (Fig. 4E, F). Critically, this proliferation was enriched in antigen-experienced populations: activated CD4^+^CD44^+^ and CD8^+^CD44^+^ phenotypes increased 3.4-fold and 3-fold, respectively (Fig. 4G, H). Selective expansion of antigen-experienced over naive T cells in the TDLN is the hallmark of clonal proliferation driven by specific antigen encounter, here most plausibly tumor-derived antigens released following mechanical disruption of the primary tumor. Functional synergy between these expanded helper and effector populations optimizes the systemic response to ensure durable protection.

Within 48 hours of a single US-NB treatment, localized mechanical disruption had initiated a coordinated systemic immune response. The expansion of antigen-experienced CD44^+^ T cell clones across both the TDLN and spleen establishes the cellular foundation for durable systemic immunity, providing long-term resistance against the recurrence and metastatic dissemination that are the primary drivers of mortality in breast cancer.

### US-NB achieves durable tumor regression and establishes systemic immune memory

Since US-NB stimulated systemic immune priming, we next assessed whether this could be translated into durable antitumor immunity. US-NB was administered three times (Days 0, 2, and 7), with the treatment schedule informed by the immune kinetics characterized in Figures 2–4. On Day 2, ultrasound application was delayed 15 minutes post-injection to allow extended intratumoral NB distribution through the partially remodeled ECM and to account for the dynamic cellular trafficking initiated by the Day 0 treatment prior to cavitation activation. A third treatment was administered on Day 7, timed to coincide with ECM structural recovery between treatments. Mice were monitored daily for body weight, caliper-based tumor measurements, and overall health (Supplementary Figure S3).

All US-NB treated mice responded to treatment. By Day 15, 85% achieved complete tumor clearance. The remaining 15% showed substantial tumor regression, surviving at least one month beyond the median endpoint of untreated controls before reaching terminal burden (Fig. 5A-B). Treated animals showed no significant or lasting weight loss throughout the study (Fig. S3).

**Figure 5.**
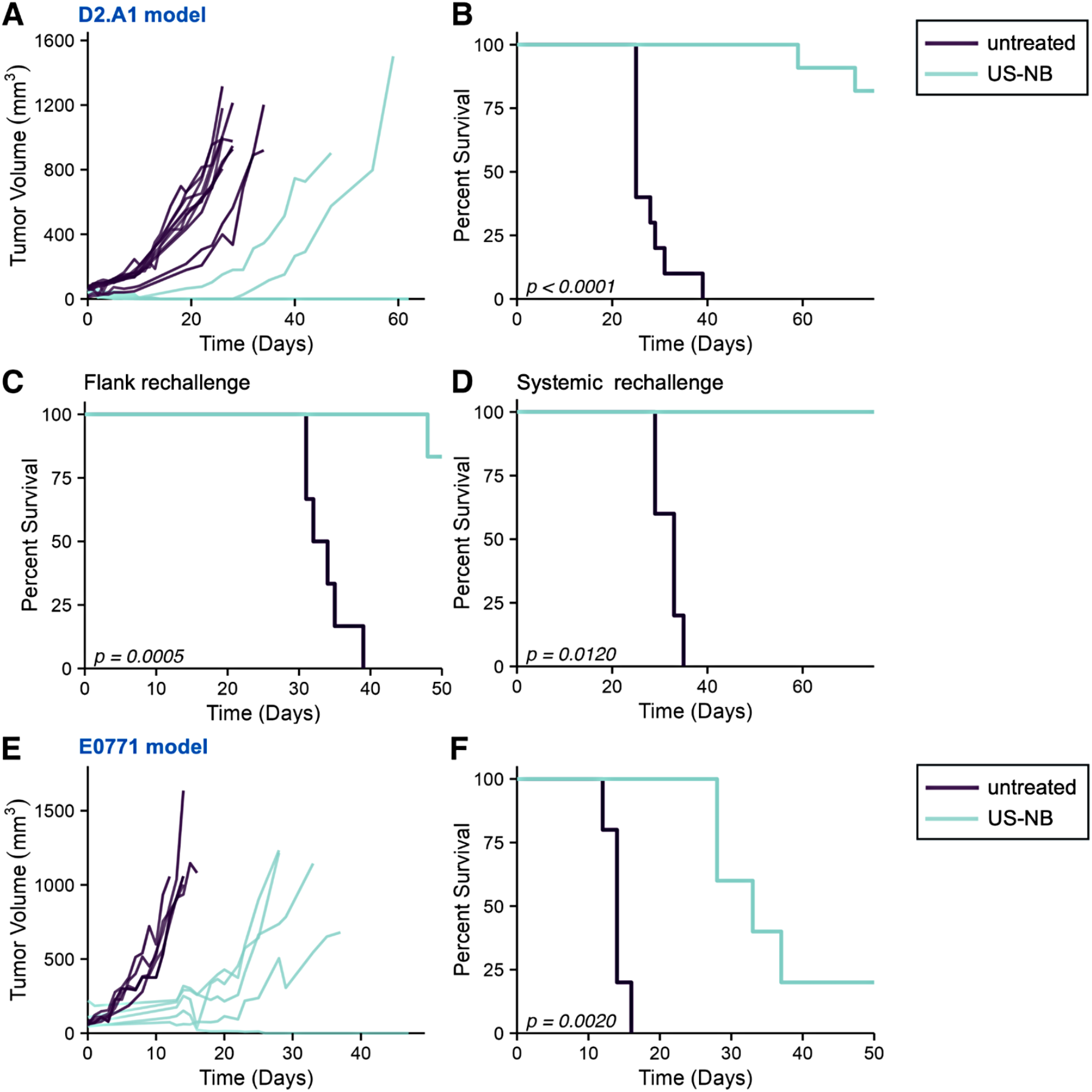
Tumor growth, survival, and rechallenge outcomes following US-NB treatment. D2A1 and E0771.LMB tumor-bearing mice received US-NB treatment on Days 0, 2, and 7. (A) Primary D2A1 tumor growth curves; volume calculated as 0.5 × length × width^2^. (B) Kaplan-Meier survival, primary D2A1 treatment (n=8 per group). (C) Kaplan-Meier survival following contralateral flank rechallenge at Day 60 post-inoculation (n=5 cured mice). (D) Kaplan-Meier survival following intravenous rechallenge (n=3 cured mice). (E) Primary E0771.LMB tumor growth curves (n=5 per group). (F) Kaplan-Meier survival, E0771.LMB primary treatment. Statistics: log-rank (Mantel-Cox) test.

To determine whether tumor clearance reflected durable immune memory, cured mice were rechallenged 60 days after initial inoculation with the same tumor inoculant on the contralateral flank. All previously treated animals mounted an immune response: 83% rejected the secondary tumor entirely, and 17% showed significantly delayed progression, reaching terminal burden approximately 20 days later than untreated controls (Fig. 5C). Resistance to a tumor implanted two months later at a distant site confirms the generation of durable circulating immune memory.

Since metastatic spread remains the primary driver of mortality in breast cancer, we evaluated whether this memory extended to circulating tumor cells. Cured mice received an intravenous inoculation of the same tumor burden used for the primary experiment and all survived without detectable disease (Fig. 5D), demonstrating that US-NB-primed systemic immunity was sufficient to control circulating tumor cells.

To evaluate whether therapeutic activity extended beyond the D2A1 TNBC background, we applied the identical protocol to E0771.LMB mammary tumor-bearing mice, a luminal B-like syngeneic model with distinct receptor expression, growth kinetics, and immune composition. As the protocol was developed and optimized in D2A1, the E0771.LMB arm was treated without modification. US-NB treatment significantly inhibited tumor growth and extended survival to at least double that of untreated controls (p=0.0020), with 20% achieving complete responses (Fig. 5E-F). These results suggest that the therapeutic activity of US-NB mechanical immunomodulation may extend across breast cancer subtypes, consistent with a mechanism that acts through biophysical tissue properties rather than tumor-specific molecular targets.

## DISCUSSION

US-NB treatment reprogrammed an immunologically cold breast tumor into one capable of mounting a durable antitumor immune response.^32^ In established TNBC tumors in mice, treatment achieved 85% complete tumor clearance and generated systemic immune memory sufficient to reject both local and circulating tumor rechallenge months later. This outcome was drug-free, required no prior tumor characterization, and was driven by the biophysical properties of tumor tissue, consistent with a mechanism that does not require specific molecular targets. By mapping the spatiotemporal distribution of NBs, the acute and sustained immune response, and the downstream therapeutic outcome, we reveal a stepwise mechanistic sequence in which mechanical disruption initiates a rapid cold-to-hot conversion. Specifically, an early reversal of myeloid immunosuppression was followed by coordinated expansion of antigen-experienced effector T cells, culminating in durable systemic immunity.

Ultrasound-activated microbubbles and nanobubbles have primarily been used to enhance the intratumoral delivery of co-administered therapeutics, including checkpoint antibodies, cytokine plasmids, and TLR agonists, where immune engagement depends on the pharmacological activity of the delivered agent rather than the mechanical stimulus itself.^33^ At the other extreme, ablative ultrasound approaches have been used to directly destroy tumor tissue. Thermal ablation strategies such as HIFU directly ablate tumor cells, but coagulative necrosis damages tissue architecture and denatures proteins,^34^ compromising the antigen integrity needed for productive immune engagement.^35^

Histotripsy, a non-thermal alternative, avoids protein denaturation by nucleating cavitation from dissolved tissue gases at peak negative pressures of 15–30 MPa, but the pressures involved reduce the entire treated volume to acellular debris, eliminating the in situ stromal scaffold on which antigen presentation and immune cell trafficking depend.^15,16^ Our approach is mechanistically distinct from all three. NB shell deformability and nanoscale dimensions enable uniform distribution throughout the entire tumor volume, unlike microbubbles which remain confined to the vascular and near-perivascular space.^17^ The mechanical forces generated by NB cavitation at ∼0.25 MPa, approximately two orders of magnitude below histotripsy thresholds, remodel the ECM without bulk cellular destruction, reducing tumor stiffness and spatial heterogeneity while preserving cell viability, as we have previously demonstrated using the same NB formulation and core acoustic parameters.^17^ The downstream immune response, including systemic T cell expansion and durable rechallenge protection, reflects a treatment that restructures the physical microenvironment to enable endogenous immune engagement independent of any pharmacological agent.

The link between local disruption and systemic immunity is evidenced by the rapid and sequential remodeling of the tumor microenvironment across both innate and adaptive compartments. mMDSC depletion was the earliest observed change, occurring within 3 hours of treatment and persisting through 48 hours. By 48 hours, antigen-presenting myeloid populations had expanded in both the spleen and tumor-draining lymph node, indicating that the immunogenic signals generated by US-NB propagated beyond the primary tumor to systemic lymphoid sites. This innate reprogramming preceded and likely facilitated the expansion of CD44^+^ effector-memory T cells: mMDSC depletion removed a primary barrier to T cell function, while macrophage polarization toward antigen-presenting phenotypes created a TME that supported T cell activation and survival.^36-39^ The upregulation of CD44^+^ antigen-experienced T cells is consistent with antigen-driven activation, linking directly to the HMGB1-mediated danger signaling and granzyme B cytotoxic activity quantified in parallel. The trajectory from an immunosuppressive (CD25^+^-dominated) to an effector and memory-forming (CD44^+^-dominated) state occurred concurrently with innate immune polarization, demonstrating a coordinated and atypically rapid innate-to-adaptive handover within 48 hours.^40,41^ Collectively, these observations support a mechanically initiated cascade that progressively dismantles the suppressive myeloid landscape of the TME.

In a prior study, we established that ultrasound alone (without NBs) applied at the same core acoustic parameters (3.3 MHz, 2.2 W/cm^2^, 50% duty cycle) produced no detectable tissue disruption or immune cell engagement in syngeneic murine breast cancer.^17^ While the present study employs a longer treatment duration to maximize cavitation energy for standalone immunomodulation, the absence of thermal elevation and the established requirement for cavitation nuclei at these acoustic intensities support the specificity of the observed effects to NB-mediated cavitation. Future work will include duration-matched US-only controls to provide formal confirmation and will optimize acoustic parameters for tumors with varying mechanical properties.

Clinically, this biophysical mechanism offers a distinct advantage in the context of aggressive breast cancers like TNBC, which is characterized by profound inter- and intra-patient heterogeneity.^42,43^ The biological landscape of metastatic lesions often diverges from the primary tumor, leading to unpredictable responses to receptor-targeted therapies or checkpoint inhibitors. Pharmacological agents are further limited by acquired resistance mechanisms such as antigen loss or multidrug resistance pathway upregulation.^44-46^ In contrast, mechanical disruption targets biophysical rather than molecular properties of tumor tissue, because a cell’s susceptibility to physical perturbation does not depend on its mutational burden or receptor profile. This positions US-NB as a candidate strategy for molecularly heterogeneous tumors that might otherwise escape pharmacological interventions, though validation beyond breast cancer subtypes is needed. The approach also offers a favorable translational profile. Ultrasound is already widely and safely used in the clinic, and the nanobubbles are composed of biocompatible phospholipids loaded with inert perfluorocarbon gas. Since the cavitation effect can be spatially directed by the transducer, treatment can be confined to tumor-bearing regions while sparing healthy tissue.

In this work, we employed intratumoral injection of NBs. This administration route limits the current approach to tumors that are readily accessible for biopsy. IT administration is not feasible for inaccessible, deep-tissue lesions or micrometastases.^47,48^ Future work will evaluate systemically delivered NBs and the ability of US-NB to modulate immunity at distant metastatic sites. The translation of these findings to clinical practice will also require adaptation of the ultrasound parameters, which were tuned to the specific dimensions and biology of murine breast cancer models. Transitioning to larger tumor volumes, different tissue contexts, or other species will require adjusting cavitation settings to generate a comparable biological effect.^49^ Future work will define these scaling parameters and further characterize the specific cytokine pathways that sustain the long-term memory phase.

In conclusion, US-NB mechanical immunomodulation converted an immunologically cold tumor into a site of active, durable antitumor immunity without pharmacological intervention. By acting on biophysical tissue properties shared across breast cancer subtypes rather than tumor-specific molecular targets, this approach circumvents the resistance pathways that typically limit immunotherapies. These data position US-NB as a drug-free strategy capable of generating systemic protective immunity from a single treatment site, with translational potential for the control of metastatic disease.

## METHODS

### Animal Studies

All animal studies were performed under an approved protocol (protocol 2016-0115) by the Institutional Animal Care and Use Committee (IACUC) of Case Western Reserve University. D2A1-FLuc-GFP cells were a gift from the Schiemann Lab (School of Medicine, Case Western Reserve University). Tumor cells were cultured in RPMI supplemented with 10% FBS and 1% PS (Gibco, Evansville, IN, USA). A small volume (30 μL) of 250,000 cancer cells was intradermally inoculated on the right flank of 6-8-week BALB/c mice (Jackson Labs, Bar Harbor, ME, USA). Weight and tumor volumes were calipered daily. Tumor volumes were calculated using a standard formula (length x width^2^ x 0.5), where length is the longest axis, and the width is perpendicular. Studies were initiated when the tumors reached a volume of 40-60 mm^3^. When tumors reached a volume of 1000 mm^3^ or mice lost 20% of their initial body weight, mice were euthanized. Survival studies were conducted by applying TUS immediately after NB injection on day 0 and 7 after initial treatment. Additionally, a single treatment was given on day 2 where TUS was applied 15 minutes after injection. Local rechallenge studies were conducted by re-inoculating mice with 250,000 tumor cells on the opposite flank 60 days after initial inoculation. Systemic rechallenge studies were conducted by re-inoculating mice with 250,000 tumor cells intravenously, 30 days after local rechallenge.

### Synthesis of nanobubbles

NBs were formulated following an established protocol.^17,50^ Briefly, DBPC, DPPA, DPPE, and mPEG2000-DSPE were dissolved in propylene glycol (PG). The lipid solution was then emulsified in PBS supplemented with glycerol. Next, 1 mL of the lipid emulsion was aliquoted into a 3-mL vial, the air inside was removed and replaced with C_3_F_8_ gas. The NBs were activated by shaking the vial using a VialMix shaker (Bristol-Myers Squibb Medical Imaging, Inc., N. Billerica, MA, USA) for 45 s, which drove bubble self-assembly. NBs were isolated from larger bubbles using centrifugation (50 rcf for 5 min; vial inverted). To prepare fluorescently labeled NBs, the same method was followed with the addition of DSPE-PEG2000-Cy5, first prepared as a lipid film through chloroform evaporation.

### Characterization of nanobubbles

Resonance mass measurement (Archimedes, Malvern Analytical Inc., Westborough, MA, USA) was used to measure the size distribution and concentration of buoyant particles (bubbles) and non-buoyant particles (lipid aggregations, micelles) with a calibrated nano-sensor (50 nm - 1 µm). The sensors were pre-calibrated using NIST-traceable 565 nm polystyrene bead standards (ThermoFisher 4010S, Waltham, MA, USA).^51^ NBs were diluted in PBS (1:1000 dilution) with at least 500 particles being measured per run (n=3 runs). Dynamic light scattering (DLS) was used to further analyze the hydrodynamic diameter (Litesizer™ 500, Anton Paar, Ashland, VA, USA). NBs underwent quality control by assessing their in vitro acoustic properties, including initial contrast enhancement and stability under ultrasound, using a tissue-mimicking agarose phantom.^51^

### Contrast Enhanced Ultrasound Imaging

Nanobubbles were administered intratumorally using a 29G insulin syringe (Exel International, Salaberry-de-Valleyfield, Quebec J6T 0E3, Canada). D2A1 tumor-bearing mice were imaged using the AplioXG SSA-790A (Toshiba Clinical Medical-Imaging Systems, Otawara-Shi, Japan) with an 18L7 probe. Mice (n=3) were imaged at 0, 90, 180 min at 3 planes per organ. Non-linear contrast signals were quantified using ImageJ and then averaged across technical replicates.

### Ex vivo Imaging

Fluorescently labeled Cy5 nanobubbles were injected intratumorally in D2A1 tumor-bearing mice and assessed after 3 hours. The tumor, liver, lungs, kidney, spleen, heart, tumor-draining lymph node and plasma were harvested. Images were taken using the IVIS (Perkin-Elmer) with the appropriate spectral unmixing and positive controls. ROIs were analyzed using Living-Image 4.3.1 for the average radiance efficiency.

### Therapeutic ultrasound

NB cavitation studies were carried out when the tumor size was ∼50 mm^3^. NBs were diluted 1:5 in PBS and ∼1x10^9^ NBs in 20 μl were intratumorally administered using a 29G insulin syringe (Exel International, Salaberry-de-Valleyfield, Quebec J6T 0E3, Canada). Therapeutic ultrasound (TUS) was applied on the tumor using an unfocused transducer with a 1 cm^2^ effective radiating area (Sonicator 740, Mettler, Anaheim, CA, USA) at 3.3 MHz, 2.2 W/cm^2^, and a 50% duty cycle for 10 min, using acoustic coupling gel. TUS parameters included a pulse repetition frequency (PRF) of 100 Hz, a pulse length of 10 ms, and an estimated peak negative pressure (PNP) amplitude of 0.25 MPa.

### Flow cytometry

Flow cytometry studies were conducted by harvesting the tumor, spleen and tumor-draining lymph node. Tumors were pre-digested in Liberase (0.4 mg/mL, Roche, IN, USA) and DNAse (0.2 mg/mL, Roche) and processed through 70 µm nylon strainers like the spleen and lymph node. These single cell suspensions were plated and then blocked with CD16/CD32 Fc block (Biolegend, San Diego, CA, USA). Cells then underwent immunophenotyping with fluorescent antibodies staining (Supplementary Table S1). For viability assessment, cells were counterstained with DAPI (BD Biosciences, Franklin Lakes, NJ, USA). Cellular ROS (Abcam) was used to stain single-cell suspensions from the tumor for DCFDA. All cells were read with the BD-LSR Fortessa and analyzed using FlowJo.

### Measurement of cytokine, chemokine and DAMPs

Tumors were submerged in 1 mL PBS and homogenized with a tissue homogenizer and then stored in -80°C. Samples underwent 3 freeze-thaw cycles and were centrifuged before storage in -80°C. ELISA was used to quantify HMGB1 and Granzyme B (CusaBio, Houston, TX, USA) in tumor tissues following the manufacturer’s instructions and absorbance readings were measured using a plate reader (Biotek, Winooski, VA, USA).

### Statistics

Statistics were performed using GraphPad Prism and data is represented as mean ± SD. Unpaired t-tests and one-way analysis of variance (ANOVA) were performed with the appropriate post-hoc tests. Multiple comparisons were conducted by comparing each group to each other and p < 0.05 was considered significant.

## ACKNOWLEDGEMENTS

This work was supported by grants from the National Institute of Biomedical Imaging and Bioengineering (R01EB028144, AAE), National Cancer Institute (R01CA253627, R01CA278633, E.K.), and pilot funding from the Case Comprehensive Cancer Center Support Grant (P30CA043703, AAE and EK). A.B. was supported by a fellowship from the NIH Interdisciplinary Biomedical Imaging Training Program (T32EB007509). I.M.H. received funding from an NIH Institutional Training Grant (T32GM007250). We acknowledge the help of Taylor J. Moon and Andrew Choi with the flow cytometry studies. We also acknowledge the Imaging Core, and Cytometry and Imaging Microscopy Core. Figure illustrations were made using R. Statistical analyses were conducted using GraphPad Prism and R. Graphical abstract was made using Nano Banana.

