## Supplementary Material for "Ultrasound-Activated Nanobubbles Induce Durable Systemic Antitumor Immunity"

### Contents

**Table S1.** Flow phenotyping for immune cell populations

**Figure S1.** Characterization of Cy5-conjugated nanobubbles

**Figure S2.** Flow cytometry gating strategy for all immunophenotyping analyses.

**Figure S3.** Body weight monitoring during US-NB treatment.

| Cell population | Immunophenotype |
|---|---|
| Pan-immune cells | DAPI-/CD45+ |
| Dendritic cells | DAPI-/CD11b+/CD11c+ |
| Macrophages | DAPI-/CD11b+/F4-80+ |
| mMDSCs | DAPI-/CD11b+/Ly6chiLy6G- |
| T cells | DAPI-/CD3e+/CD4+ or CD8+ |
| NK cells | DAPI-/CD3e-/CD49b |
| Neutrophils | DAPI-/CD11b+/Ly6G+CD244- |
| Activated and memory T cells | DAPI-/CD3e+/CD4+/CD44+ or DAPI-/CD3e+/CD8+/CD44+ |
| Activated or regulatory T cells | DAPI-/CD3e+/CD4+/CD25+ or DAPI-/CD3e+/CD8+/CD25+ |
| Antigen presenting macrophages | DAPI-/CD11b+/F4-80+/MHC2+ |
| M1 Macrophages | DAPI-/CD11b+/F4-80+/MHC2+/CD86+ |
| M2 Macrophages | DAPI-/CD11b+/F4-80+/CD206+ |

**Supplementary Table S1.** Flow phenotyping for immune cell populations

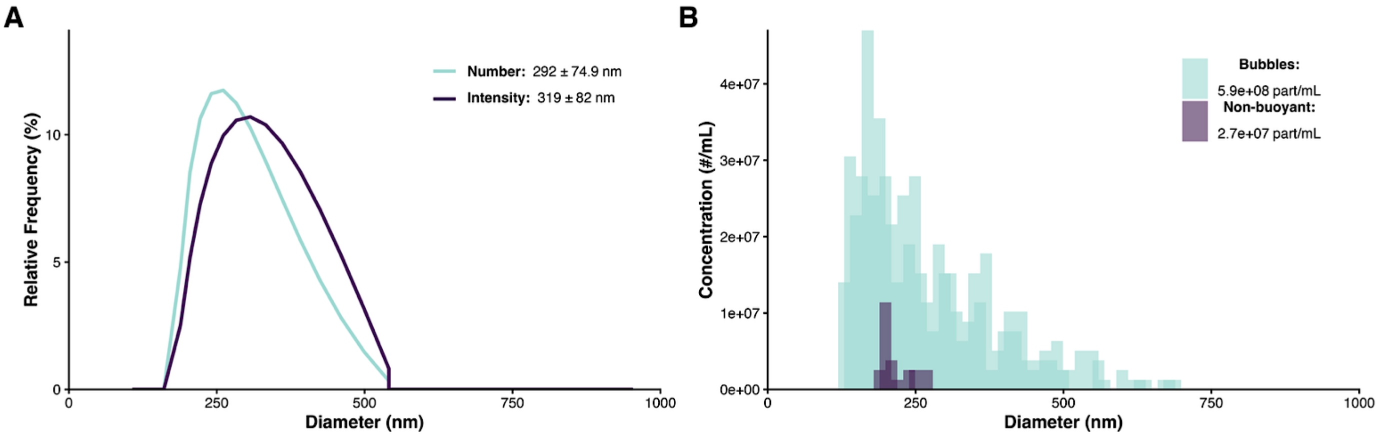


**Supplementary Figure S1. Characterization of Cy5-conjugated nanobubbles**. **(A)** Hydrodynamic size distribution of Cy5-NBs measured by dynamic light scattering (DLS; Litesizer 500, Anton Paar), reported as number-weighted (292 ± 74.9 nm) and intensity-weighted (319 ± 82 nm) distributions. **(B)** Buoyant particle concentration measured by resonant mass measurement (RMM; Archimedes, Malvern; 100 nm to 2 μm range; n=3 runs, at least 500 particles per run) at a 1:1000 dilution in PBS, yielding an undiluted buoyant NB concentration of 5. 9 × $10^{11}$ particles/mL and a non-buoyant fraction of 2.7 × $10^{10}$ particles/mL.

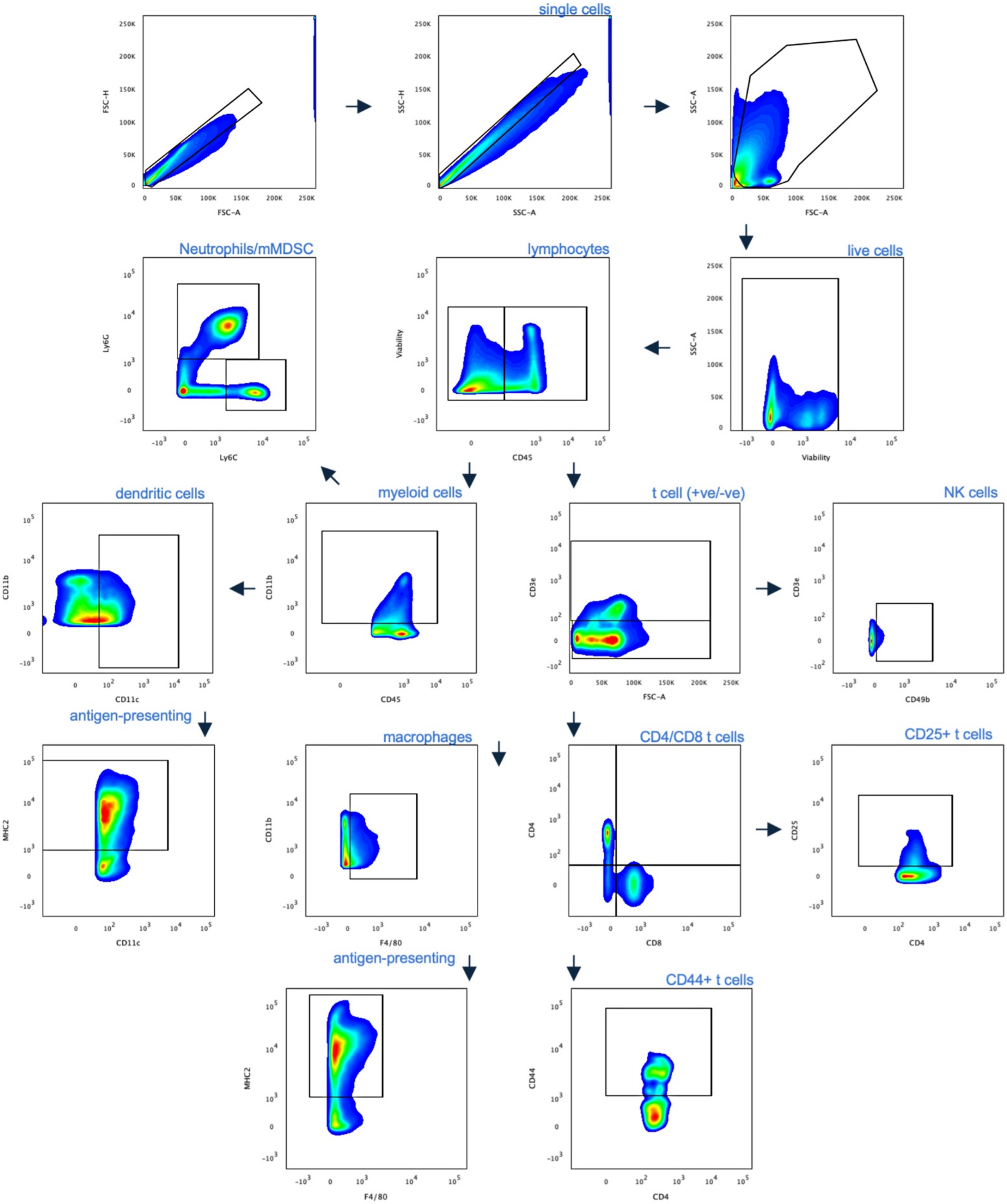


**Supplementary Figure S2. Flow cytometry gating strategy for all immunophenotyping analyses.** Representative hierarchical gating plots applied to single-cell suspensions from D2A1 tumors, tumor-draining lymph nodes, and spleens. All immune populations quantified in this study are defined in the accompanying table. Representative plots shown from a D2A1 tumor-bearing mouse.

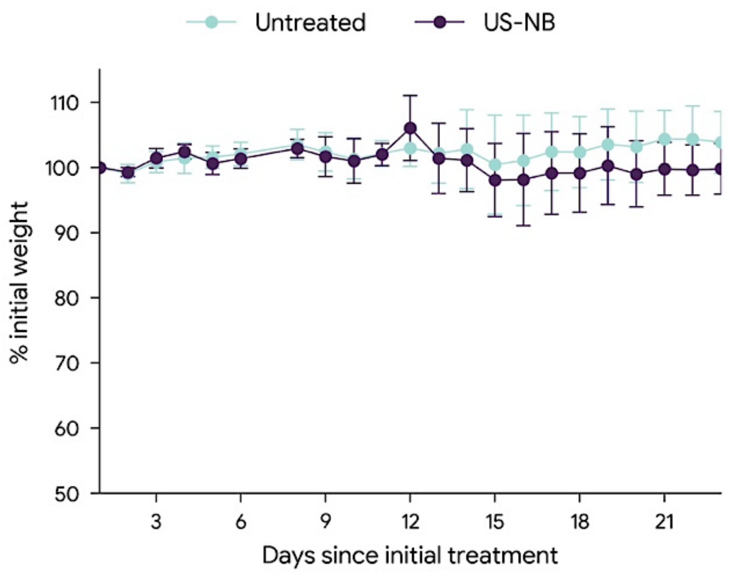


**Supplementary Figure S3. Body weight monitoring during US-NB treatment.** Body weight of D2A1 tumor-bearing mice in the US-NB and untreated groups, recorded daily from the day of initial treatment. Data expressed as percentage of initial body weight at Day 0 (n=11 per group; mean ± SD). No significant or lasting weight loss was observed in US-NB treated animals throughout the study period.
